# Age-Related Upregulation and Strong Expression Correlation of the Proinflammatory Cytokines IL-1β and IL-6 Across Multiple Segments of the Mouse Eye

**DOI:** 10.64898/2026.01.09.698735

**Authors:** Haley Greene, Irina Sbornova, Michy P. Kelly

**Author notes:** Corresponding author, University of Maryland School of Medicine 20 Penn St, HSFII Rm S218e, Baltimore, MD 21201. contributed equally (Haley Greene, Irina Sbornova).

## Abstract

Emerging evidence suggests that ocular inflammation increases with age and is associated with various disease states. That said, the majority of studies suffer from a significant limitation—they only focus on a single segment of the eye. This represents a limitation because age-related increases in neuroinflammatory markers are not necessarily uniform within an organ, and other age-related changes in the eye are known to occur in a segment-specific manner. The present study aims to address this gap by comparting/contrasting age-related changes in the proinflammatory cytokines IL-6 and IL-1β across multiple mouse ocular segments: the (i) anterior segment (i.e., cornea, ciliary body and muscle, and zonules), (ii) retina, and (iii) posterior segment (i.e., sclera, choroid, Bruch’s membrane, retinal pigmented epithelium, and parts of the optic nerve). IL-6 and IL-1β were selected as targets since they exhibit differential regional patterns of age-related increases within the brain. Eyes were collected on postnatal days (P) P28, P56, P98, P200 and P500 and processed by Western blot. Both IL-6 and IL-1β protein levels increased across the lifespan in all three eye segments. Interestingly, correlational analyses revealed that IL-1β and IL-6 expression correlated with each other not only within individual eye segments, but also across segments. In old mice, IL-1β and IL-6 levels also correlated with expression of phosphodiesterase 11A (PDE11A), an enzyme known to regulate neuroinflammation. Together, these findings suggest that inflammaging in the eye is broadly controlled by a systemic governor, unlike the brain that shows region-specific changes in cytokines with age.

## INTRODUCTION

Aging is accompanied by a persistent, low-grade inflammatory state known as inflammaging [1–6]. Inflammaging occurs due to activation of immune cells, which ultimately increase levels of proinflammatory cytokines and decrease levels of anti-inflammatory cytokines [7–18]. This process is associated with increased mortality [19–26] and age-related disease [27–34]. Thus, it is critical to gain a clear understanding of when and where inflammation arises within the body. Notably, several studies suggest that age-related increases in neuroinflammatory markers are not uniform across all organs, nor even within an organ [17, 18, 35–37]. For example, in the brain, we previously demonstrated that age-related increases in IL-6 protein expression occurred in the mouse hippocampus, prefrontal cortex, striatum, and cerebellum, whereas age-related increases in IL-1β protein only occurred in the hippocampus [37]. However, a critical unanswered question is how age-related inflammation is organized within complex organs that contain multiple anatomically and functionally distinct compartments.

Emerging evidence suggests that ocular inflammation increases with age and is associated with various disease states of the eye [38–64]. For example, age-related increases in protein expression of the proinflammatory cytokine IL-6 have been reported in the retina [48] and aqueous humor [49], while age-related increases in the proinflammatory cytokine IL-1β have been noted in the aqueous humor, conjunctiva, lacrimal glands, and tears [40, 47, 49, 60]. Interestingly, elevated retinal levels of IL-6 and IL-1β along with tumor necrosis factor α (TNFα) and transforming growth factor- β2 (TGF-β2) were associated with glaucoma [44]. Age-related increases in proinflammatory macrophages and/or microglia in the retina were also associated with glaucoma [44] along with retinal degeneration [46]. In contrast, wet age-related macular degeneration (wet AMD) was associated with increased levels of the inflammatory chemokines CXCL10, CCL14, CXCL16, CXCL7, and CCL22 in the aqueous humor of patients versus controls [45]. Finally, the development of dry eye disease has been associated with age-related increases in TNF in the lacrimal gland, which drives goblet cell loss and ectopic lymphoid structure formation [40]. Together, these findings suggest that inflammatory markers may represent biomarkers and/or therapeutic targets for the stratification, diagnosis or treatment of age-related eye diseases. Notably, these associations have largely been characterized within individual ocular compartments, leaving open the question of whether inflammatory changes generalize across the eye or remain segment-specific.

Such analyses are at a disadvantage since other age-related physiological changes in the eye occur in a segment-specific manner. This stands in contrast to the brain, where regional heterogeneity in age-related cytokine regulation is well established, raising the possibility that inflammaging in the eye could follow either compartment-specific or organ-wide organizational principles. For example, we previously showed that protein expression of the enzyme PDE11A4, which regulates neuroinflammation [65–67], changes across the lifespan in an eye-segment specific manner, with increases in the anterior segment and decreases in the retina [68]. These prior findings suggested that, like other age-related processes in the eye, inflammatory signaling might also be differentially regulated across ocular segments. Similarly, metabolic markers exhibit much more pronounced age-related changes in the retina and optic nerve versus the cornea, choroid, and lens [69].

Together, these results suggest that inflammatory changes identified in one segment of the eye may not generalize to other segments, and may even follow opposite patterns. To address this gap, the present study pursued three objectives. First, we quantified age-related changes in the proinflammatory cytokines IL-6 and IL-1β across multiple ocular segments, including the anterior segment, retina, and posterior segment. Second, we tested whether cytokine expression levels were correlated within and across these segments, providing insight into potential organizational principles governing ocular inflammaging. Finally, we examined the relationship between cytokine expression and levels of the neuroinflammation-regulating enzyme PDE11A4 to explore potential modulatory associations. We show here that IL-6 and IL-1β protein levels increase across the lifespan in all three segments of the eye, with the extent of these changes correlated both within and between individual eye segments.

## METHODS

### Subjects

Experiments herein employed eye samples processed for Western blotting in a previous study characterizing ocular PDE11A4 protein expression levels [70]. Since the original goal for these samples focused on PDE11A4, eyes were harvested from the C57BL/6NCrl-PDE11A^em1(IMPC)Mbp/Mmucd^ knock-out mouse line from the Mutant Mouse Resource and Research Center (donating investigator: Kent Lloyd, University of California). Here we use samples only from the wild-type mice. As described in our previous publication, the mice were socially housed and had access to water and food ad libitum, with procedures approved by the ethical committee of the Royal Netherlands Academy of Arts and Sciences (KNAW, Amsterdam, The Netherlands) and care of the animals performed in compliance with the ARVO Statement for the Use of Animals in Ophthalmic and Vision Research, as well as the European Communities Council Directive 2010/63/EU.

### Tissue harvest

As previously described [70], the mice were killed by cervical dislocation with isoflurane anesthesia. The eyes were harvested fresh and dissected in phosphate-buffered saline on wet ice. The ocular tissue was separately isolated for the (1) anterior segment (i.e., cornea, ciliary body and muscle, and zonules), (2) retina, and (3) posterior segment (i.e., sclera, choroid, Bruch’s membrane, retinal pigmented epithelium, and parts of the optic nerve). The tissue was pooled for both eyes of the same mouse, put on dry ice, and kept at −80 degrees until further processing. All the tissue samples were isolated between 8 a.m. and 2 p.m.

### Preparation of samples for Western blots

As previously described [70], all tissue samples were stored at −80 °C and kept on dry ice until processed. For analysis of individual eye segments, samples were sonicated for 3 cycles using ice-cold buffer [20 mM Tris-HCl/2 mM MgCl2/0.5% Triton X-100 with protease inhibitor (Pierce #A32953) and phosphatase inhibitor cocktail 3 (#P0044 Sigma ST Louis, MO USA)]. The homogenized retina samples were precleared by centrifuging at 4 °C for 10 min at 1000× *g* and transferring the resultant supernatant to a new test tube. Any solid tissue that remained in the anterior and posterior segment samples was shaken down to the bottom of the tube, and the liquid portion was transferred to a new test tube because preclearing greatly reduced the protein concentrations of the samples. The total protein concentration for each sample was determined using the DC Protein Assay kit (BioRad, Inc.; Hercules, CA, USA) according to the manufacturer’s protocol, and samples were prepared at 2.3-3 µg/uL using Invitrogen sample buffer (#NP0007) and sample-reducing agent (#NP0009) as per manufacturer’s protocol. The samples were then stored at −80 °C until processed by Western blot.

### Western Blotting

Westerns blots were performed as previously described in [70–73]. 11 µL of each sample was loaded onto 4–12% Bis-Tris NuPAGE polyacrylamide gradient gels (Life Technologies; Bedford, MA, USA) for electrophoresis. The protein was transferred onto a 0.45µm nitrocellulose blotting membrane (#10600008; Amersham, Buckinghamshire, United Kingdom), which was then stained with Ponceau S (#6266-79-5; Fisher Scientific, Waltham, MA USA). Ponceau S was used as a loading control based on the best practice statement of the *Journal of Biological Chemistry* [74]. The membranes were blocked in 5% milk/0.1% Tween20. The whole-eye blots were probed overnight at 4 °C with one of two cytokine antibodies: IL-6 (ARC0962; 1:2500; Life Technologies; Bedford, MA, USA) and IL-1β (M421B; 1:1000; ThermoFisher Scientific, Waltham, MA USA). The membranes were then incubated with an anti-mouse secondary antibody (115-035-146; 1:10,000; Jackson Immunoresearch,West Grove, PA USA). The protein bands were visualized using the WesternSure Premium Chemiluminescent Substrate (#926-95000 LI-COR; Lincoln, NE USA), with multiple film exposures to ensure densities fell within the linear range of the film. All protein expression was normalized to Ponceau S staining intensity as a loading control, and densitometry was conducted on films scanned in at 1200 dpi using ImageJ software v1.48 (NIH).

2.12 Statistical analysis. As previously described [70, 71], data were collected by researchers blind to treatment, and the experiments were designed to counterbalance technical variables across the biological variables. ImageJ (NIH) was used for collecting the densitometry data. Any difference in the overall expression between gels due to differences in antibody binding efficiency, film exposure duration, etc. were normalized by expressing data as a fold change of a given gel’s p98 group. Data were analyzed by Sigmaplot 11.2 (San Jose, CA, USA). Since each data set failed assumptions of normality (Shapiro–Wilk test) and equal variance (Levene’s test); therefore, non-parametric ANOVA on Ranks were used. Note that while males and females were included in each experiment, we were insufficiently powered to formally analyze for the effect of sex.

Individual data points for males are plotted as squares and females as circles to enable the reader to visually inspect the data for potential sex effects. Following a significant main effect, *post hoc* analyses were performed using the Student–Newman–Keuls or Dunn’s method. However, Sigmaplot does not yield specific p-values following non parametric tests, only ‘yes’ or ‘no’ for whether the p-value is <0.05 (our defined level of significance). Correlational analyses were conducted using Pearson Product Moment Correlations, with raw p-values corrected for multiple comparisons using a false discovery rate (FDR) calculation. Graphs were generated with GraphPad Prism 9 software. Data are plotted as mean ± SEM, with individual data points shown overtop (circles = females; squares = males).

## RESULTS

To determine if cytokine expression increased with age across eye segments in the mouse, we measured expression of the proinflammatory cytokines IL-1β and IL-6 across the anterior segment (i.e., cornea, ciliary body and muscle, and zonules), posterior segment (i.e., sclera, choroid, Bruch’s membrane, retinal pigmented epithelium, and parts of the optic nerve) and retina of male and female mice aged postnatal day (P) 28-500. Significant age-related increases in IL-1β protein expression were measured in anterior segment (H(4)=29.36, P<0.0001; Post hoc: P28 and P56 vs. P98, P200 and P500, P<0.05 each), posterior segment (H(4)=32.47, P<0.0001; Post hoc: P28 and P56 vs. P98, P200 and P500, P<0.05 each; Post hoc: P98 vs. P200 and P500, P<0.05 each) and retina (H(4)=23.74, P<0.0001; Post hoc: P28 vs. P98, P200, and P500, P<0.05 each; Posthoc: P56 vs. P500, P<0.05). Significant age-related increases in IL-6 protein expression were also measured in anterior segment (H(4)=29.48, P<0.0001; Post hoc: P28 and P56 vs. P98, P200 and P500, P<0.05 each), posterior segment (H(4)=33.46, P<0.0001; Post hoc: P28 and P56 vs. P98, P200 and P500, P<0.05 each; Post hoc: P98 vs. P200 and P500, P<0.05 each) and retina (H(4)=22.48, P=0.0002; Post hoc: P28 and P56 vs. P98, P200, and P500, P<0.05 each).

Next, we determined if IL-1β and IL-6 protein levels correlated with each other within and between eye segments. We also determined if cytokine expression levels in these samples were correlated with protein expression of PDE11A4, an enzyme previously shown to change in expression with age in these specific eye segment samples [70] and to regulate neuroinflammatory markers in the brain [65–67]. To power these analyses and determine if patterns might evolve with age, we grouped the youngest two groups (adolescent-immature adult: P28 and P56) and the oldest two groups (mature adult: P200 and P500). IL-1β and IL-6 expression were strongly correlated with each other within the anterior and posterior segments of both age groups, and the retina of the older group (Figure 3, Table). Further, IL-1β and IL-6 expression were correlated with themselves and each other across eye segments, with posterior cytokine levels correlating with those measured in the anterior segment and retina of both age groups. Finally, in older animals, anterior segment PDE11A4 protein levels were positively correlated with both cytokines within that same segment, whereas retinal PDE11A4 protein levels were negatively correlated with cytokine levels in the anterior and posterior segment. Together, these findings indicate that the variability observed in cytokine expression is unlikely to reflect random blot-to-blot technical variation and instead reflects biologically meaningful individual differences that may be related to PDE11A4 expression levels.

**Table.** Pearson Product Moment Correlations for expression of IL-1β, IL-6, and PDE11A4 protein across eye segments.

|  | Anterior |  |  | Posterior |  |  | Retina |  |  | Anterior |  |  | Posterior |  |  | Retina |  |  |
| --- | --- | --- | --- | --- | --- | --- | --- | --- | --- | --- | --- | --- | --- | --- | --- | --- | --- | --- |
| | younger | IL-6 | PDE11A4 | IL-1β | IL-6 | PDE11A4 | IL-1β | IL-6 | PDE11A4 | older | IL-6 | PDE11A4 | IL-1β | IL-6 | PDE11A4 | IL-1β | IL-6 | PDE11A4 |
| Anterior<br>IL-1β | r= | 0.981 | 0.166 | 0.927 | 0.886 | -0.060 | 0.775 | 0.493 | -0.263 | r= | 0.852 | 0.799 | 0.477 | 0.669 | -0.192 | 0.253 | 0.209 | -0.564 |
|  | raw p= | <0.001 | 0.539 | <0.001 | <0.001 | 0.826 | <0.001 | 0.052 | 0.326 | raw p= | <0.001 | <0.001 | 0.062 | 0.005 | 0.477 | 0.345 | 0.438 | 0.028 |
|  | FDR-p= | <0.001 | 0.718 | <0.001 | <0.001 | 0.959 | 0.002 | 0.145 | 0.617 | FDR-p= | 0.001 | 0.004 | 0.111 | 0.016 | 0.520 | 0.414 | 0.493 | 0.054 |
| Anterior<br>IL-6 | r= |  | 0.211 | 0.898 | 0.891 | -0.086 | 0.848 | 0.496 | -0.187 | r= |  | 0.718 | 0.644 | 0.671 | -0.303 | 0.567 | 0.435 | -0.607 |
|  | raw p= |  | 0.433 | <0.001 | <0.001 | 0.751 | <0.001 | 0.051 | 0.488 | raw p= |  | 0.003 | 0.007 | 0.004 | 0.254 | 0.022 | 0.092 | 0.016 |
|  | FDR-p= |  | 0.649 | <0.001 | <0.001 | 0.901 | <0.001 | 0.152 | 0.675 | FDR-p= |  | 0.015 | 0.023 | 0.020 | 0.351 | 0.047 | 0.150 | 0.039 |
| Anterior<br>PDE11A4 | r= |  |  | 0.041 | 0.053 | -0.279 | 0.375 | -0.058 | -0.259 | r= |  |  | 0.473 | 0.588 | -0.313 | 0.344 | 0.082 | -0.812 |
|  | raw p= |  |  | 0.881 | 0.846 | 0.295 | 0.152 | 0.832 | 0.333 | raw p= |  |  | 0.075 | 0.021 | 0.256 | 0.209 | 0.772 | <0.001 |
|  | FDR-p= |  |  | 0.933 | 0.922 | 0.591 | 0.343 | 0.936 | 0.571 | FDR-p= |  |  | 0.129 | 0.048 | 0.341 | 0.301 | 0.794 | 0.004 |
| Posterior<br>IL-1β | r= | | | | 0.815 | 0.231 | 0.645 | 0.697 | -0.221 | r= | | | | 0.832 | -0.254 | 0.674 | 0.670 | -0.644 |
|  | raw p= |  |  |  | <0.001 | 0.389 | 0.007 | 0.003 | 0.410 | raw p= |  |  |  | <0.001 | 0.343 | 0.004 | 0.004 | 0.010 |
|  | FDR-p= |  |  |  | <0.001 | 0.636 | 0.023 | 0.011 | 0.642 | FDR-p= |  |  |  | 0.001 | 0.426 | 0.022 | 0.018 | 0.027 |
| Posterior<br>IL-6 | r= |  |  |  |  | -0.032 | 0.696 | 0.483 | -0.354 | r= |  |  |  |  | -0.289 | 0.633 | 0.549 | -0.612 |
|  | raw p= |  |  |  |  | 0.906 | 0.003 | 0.058 | 0.178 | raw p= |  |  |  |  | 0.278 | 0.009 | 0.028 | 0.015 |
|  | FDR-p= |  |  |  |  | 0.932 | 0.010 | 0.150 | 0.377 | FDR-p= |  |  |  |  | 0.357 | 0.026 | 0.056 | 0.039 |
| Posterior<br>PDE11A4 | r= |  |  |  |  |  | -0.154 | 0.432 | -0.023 | r= |  |  |  |  |  | -0.233 | 0.021 | 0.357 |
|  | raw p= |  |  |  |  |  | 0.569 | 0.095 | 0.933 | raw p= |  |  |  |  |  | 0.384 | 0.938 | 0.191 |
|  | FDR-p= |  |  |  |  |  | 0.731 | 0.228 | 0.933 | FDR-p= |  |  |  |  |  | 0.446 | 0.938 | 0.286 |
| Retinal<br>IL-1β | r= | | | | | | | 0.261 | -0.097 | r= | | | | | | | 0.748 | -0.449 |
|  | raw p= |  |  |  |  |  |  | 0.329 | 0.720 | raw p= |  |  |  |  |  |  | 0.001 | 0.094 |
|  | FDR-p= |  |  |  |  |  |  | 0.592 | 0.894 | FDR-p= |  |  |  |  |  |  | 0.006 | 0.146 |
| Retinal<br>IL-6 | r= |  |  |  |  |  |  |  | -0.210 | r= |  |  |  |  |  |  |  | -0.184 |
|  | raw p= |  |  |  |  |  |  |  | 0.435 | raw p= |  |  |  |  |  |  |  | 0.513 |
|  | FDR-p= |  |  |  |  |  |  |  | 0.626 | FDR-p= |  |  |  |  |  |  |  | 0.543 |

FDR, false detection rate; younger = P28 + P56; older = P200 + P500; Bolding indicates FDR-corrected p<0.05.

**Figure 1.**
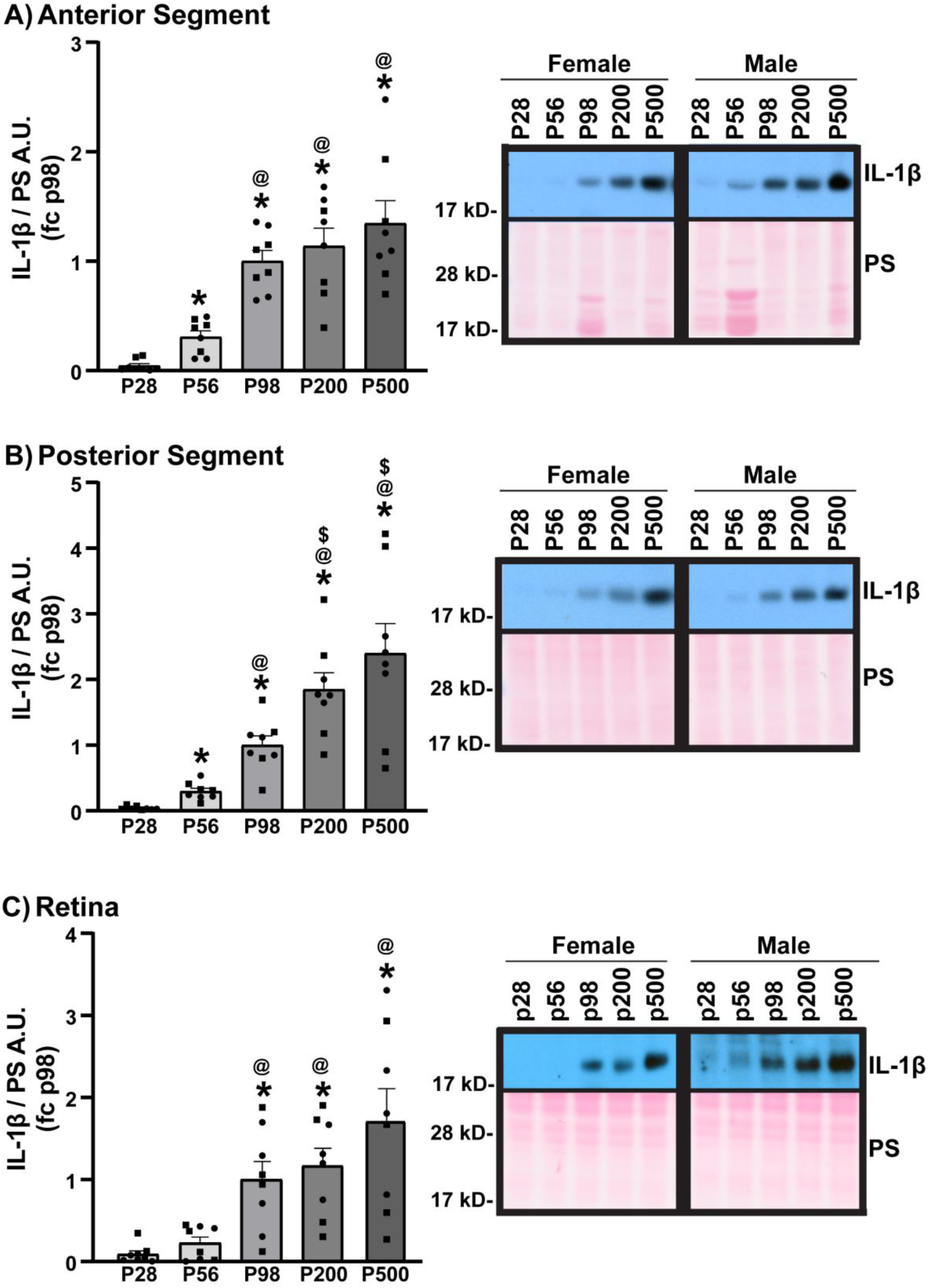
Western blots reveal age-related increases in IL-1β protein expression across all the three eye segments. Quantification of Western blots from mice aged postnatal day (P) 28-500 all demonstrate significant age-related increases in IL-1β protein expression in the A) anterior segment, B) posterior segment, and C) retina. A.U. = arbitrary units; kD = kilodalton. Data expressed as mean ±SEM with individual data points (female, circles; male, squares). *Post hoc* *vs. P28, ^@^vs. P56, ^$^vs. P98, p< 0.05.

**Figure 2.**
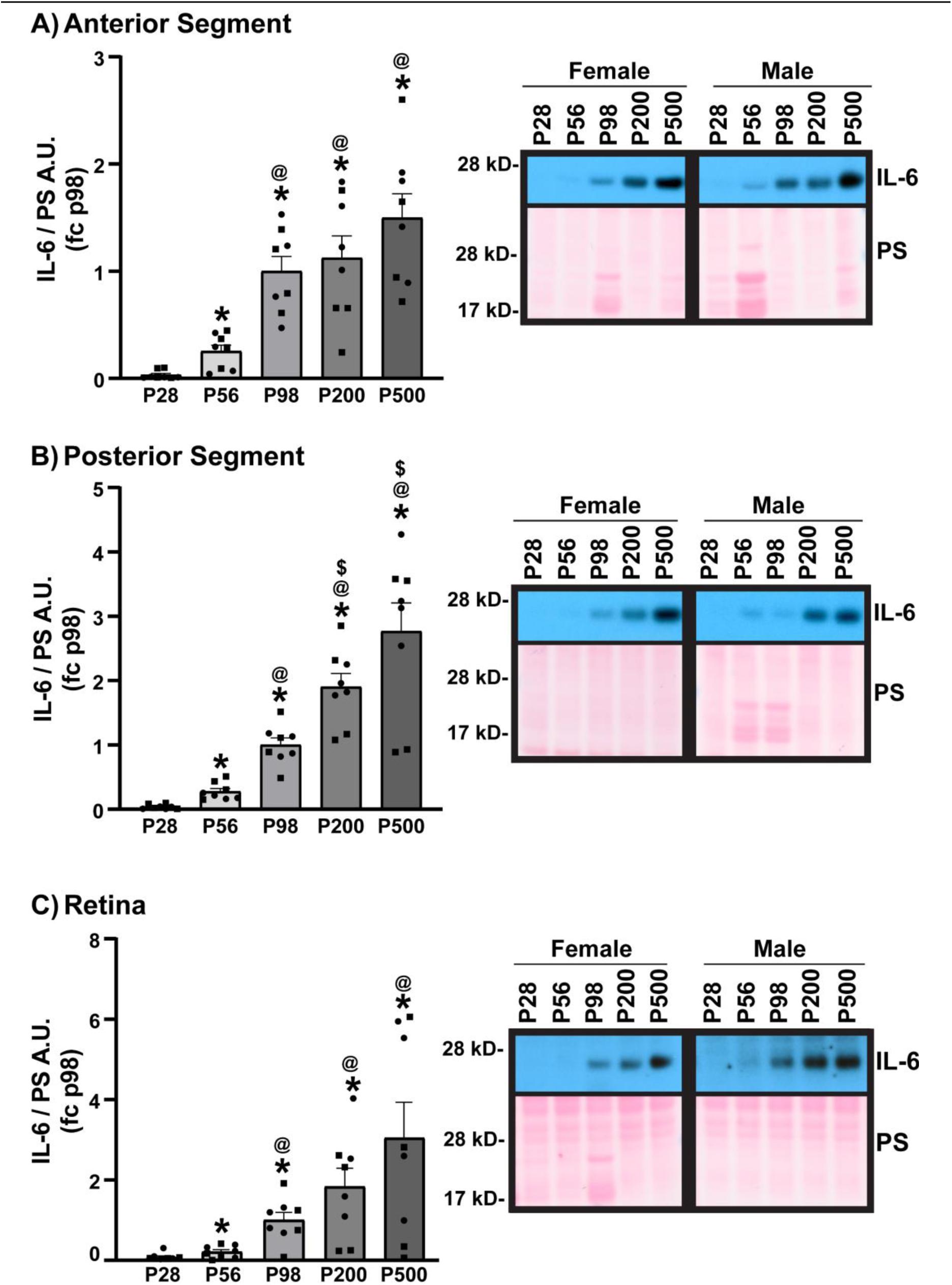
Western blots showed IL-6 protein expression also increases with age across all three eye segments. Quantification of Western blots from mice aged postnatal day (P) 28-500 all reveal significant age-related increases in IL-6 protein expression in the A) anterior segment, B) posterior segment, and C) retina. A.U. = arbitrary units; kD = kilodalton. Data expressed as mean ±SEM with individual data points (female, circles; male, squares). *Post hoc* *vs. P28, ^@^vs. P56, ^$^vs. P98, p< 0.05.

**Figure 3.**
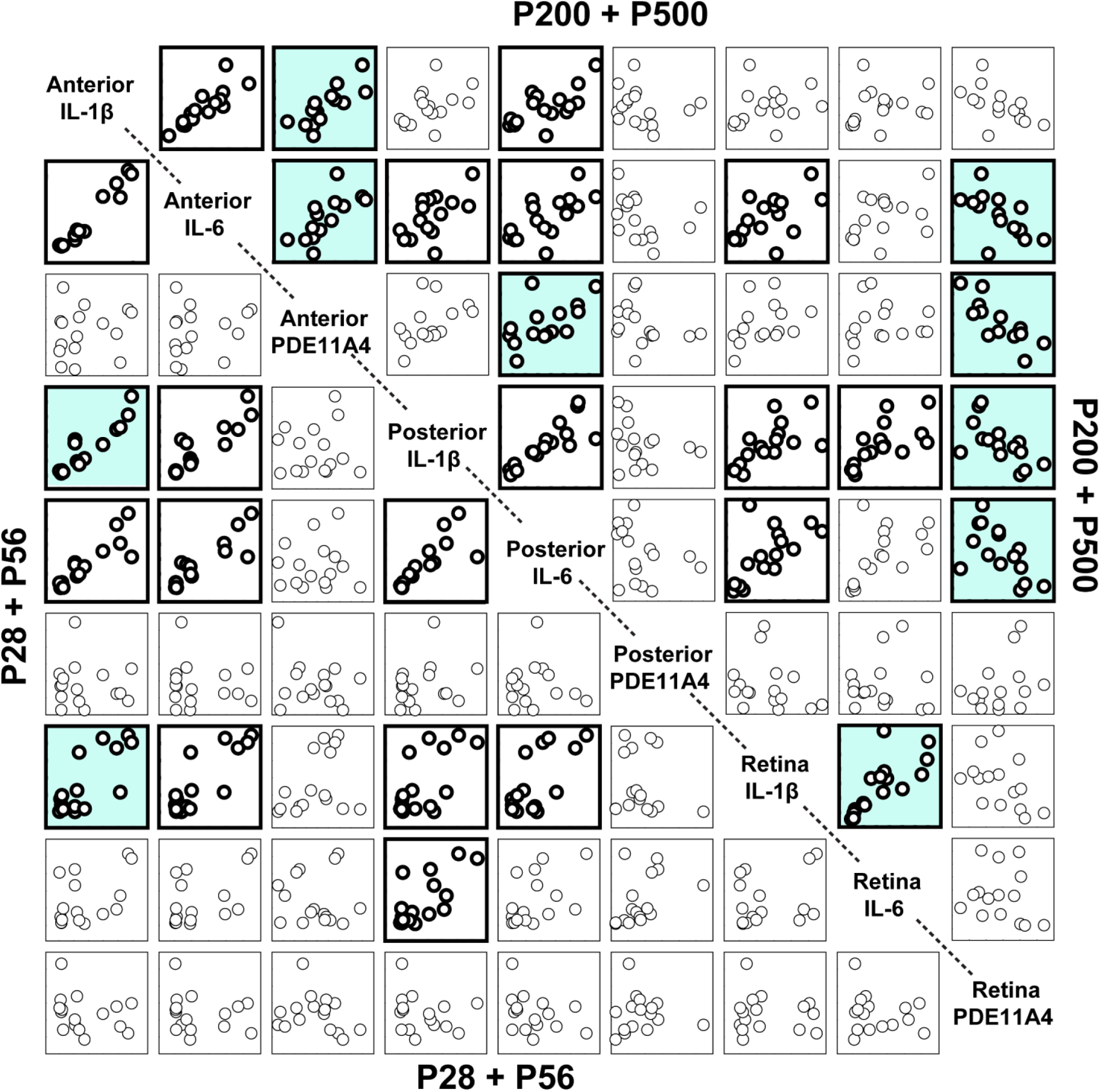
IL-1β and IL-6 protein expression correlate with each other and PDE11A4 protein expression in the older mouse groups. IL-1β and IL-6 correlate with each other within and between the anterior and posterior segments in both age groups. In contrast, retinal IL-1β and IL-6 only correlate with each other and posterior segment cytokine expression in older mice. Interestingly, anterior segment PDE114 protein expression positively correlated with anterior segment IL-1β and IL-6 in old mice; whereas, retinal PDE11A4 protein expression in the older mice correlated negatively with both anterior PDE11A4 protein expression as well as anterior and posterior segment cytokine expression. Bolding represents FDR-corrected p = 0.039 to <0.001. White background indicates correlation is significant in both age groups; colored background indicates correlation is unique to one of the age groups. See Table for r and p-values.

## DISCUSSION

The present study addressed multiple objectives aimed at characterizing both the magnitude and organization of age-related inflammatory changes in the mouse eye. With respect to our first objective, we found that the proinflammatory cytokines IL-6 and IL-1β increase with age across the retina, anterior segment, and posterior segment. To address our second objective, we examined whether expression levels of these cytokines were correlated within and across ocular segments and observed strong positive correlations both within individual segments and between distinct segments of the eye. The combination of these findings—that both cytokines increase across all ocular compartments and that their expression levels are coordinated not only within but also across segments—suggests that ocular inflammaging is organized in a globally coordinated, rather than compartment-specific, manner. This pattern stands in contrast to our previous findings in the brain, where age-related increases in IL-6 are widespread but increases in IL-1β are regionally restricted across multiple mouse and rat strains [37]. Together, these observations indicate that inflammaging may be governed by distinct organizational principles in the eye versus the brain, despite both being complex, multicompartment organs.

In addition to defining organizational patterns of ocular cytokine expression, a third objective of this study was to explore whether age-related inflammatory changes in the eye are associated with expression of phosphodiesterase 11A4 (PDE11A4), an enzyme previously shown to regulate neuroinflammatory markers in the brain [65–67]. We found that cytokine levels in older mice were significantly correlated with PDE11A4 expression in a segment-dependent manner, with positive associations observed between PDE11A4 and cytokine levels within the anterior segment, but negative associations observed between retinal PDE11A4 levels and cytokine expression in other ocular compartments. Notably, retinal PDE11A4 expression was also inversely correlated with anterior segment PDE11A4 levels. **Consistent with this opposing relationship, we previously showed that PDE11A4 expression decreases across the lifespan in the retina but increases in the anterior segment** [Sbornova et al., 2023]. Together, these findings suggest that PDE11A4-associated signaling may reflect opposing or compensatory states across ocular segments rather than uniform regulation throughout the eye. In this context, retinal PDE11A4 may index a state that is inversely related to the broader ocular cytokine milieu, rather than directly mirroring local inflammatory tone. While these findings are correlational and do not establish causality or directionality, they highlight PDE11A4 as a candidate modulator of age-related ocular inflammation and motivate future studies aimed at disentangling local versus cross-compartment influences on inflammatory signaling.

The age-related increases in ocular IL-6 and IL-1β protein observed in the present study align with a growing body of literature linking aging to heightened inflammatory signaling in the eye. A 1.5-fold increase in retinal IL-6 protein concentration was found in 8 month-old versus 3 month-old mice [48], paralleling the two-fold increase in IL-6 protein expression we observed between P98 (∼3 months) and P200 (∼7 months old) in both the retina and posterior segment. Similarly, adult humans showed a six-fold increase in IL-6 protein levels relative to children in the aqueous humor, a key immune component of the anterior segment [49]. This again is quite consistent with our observing a 3-5-fold increase in IL-6 between P56 and P98-500 in the anterior segment. IL-1β levels also increase with age between childhood and adulthood in the human aqueous humor [49], as it did in the anterior segment of our mice between P56 and P200-500. Elevated IL-1β protein and mRNA levels have also been observed in aged mouse conjunctiva, lacrimal glands, and tears [40, 47, 60] and are associated with several age-related ocular diseases such as AMD [45, 50, 51], diabetic retinopathy [52–55], and glaucoma [56–58]. Other proinflammatory cytokines similarly increase with age, including TNF-α, IL-3, IL-2, and IL-21 in human tears [59] and TNF, IL-18, IL-2, IFN-γ, IL-12p40, IL-17, and IL-10 levels in aged mouse tears [40, 47]. In addition, both IL-12 and IL-8 are increased with age in conjunctiva tissue [38, 60], and TNF-α is increased in the lacrimal glands of aged C57BL/6 mice [40, 47].

Within the field of inflammation, sex-specific differences have received substantial attention, although findings remain inconsistent. Several studies indicated that aged females exhibit a stronger inflammatory response than males [37, 75–79]. In contrast, others found higher neuroinflammation in males, particularly following immune challenges or disease states [80–83]. Here, we did not observe any clear sex differences in ocular IL-6 and IL-1β protein levels. This again contrasts with our previous study in the brain where we saw age-related increases in IL-6—but not IL-1β—were more pronounced in females than males in several brain regions across mouse strains [37]. Our findings here, thus, compliment a growing body of work that suggests sex-specific inflammatory changes are complex and highly specific to the neuroinflammatory marker measured as well as the tissue and disease state studied [37, 75–83].

While the present study may provide perspective into ocular inflammaging, several limitations should be acknowledged. First, mice were only aged to P500 (∼16.5 months old), which may limit the relevance of our findings to the later stages of aging. Second, the root cause of the cytokine increases was not explored (e.g., measurement of microglia/macrophages, the causality of PDE11A4 expression levels). Third, the functional consequences of these age-related increases in ocular cytokines were not tested. As such, it remains to be determined if the age-related increases in ocular IL-6 and IL-1β measured herein represent a pathological or protective process. Direct evidence linking proinflammatory cytokine elevations and tissue degeneration in the eye remains limited [46]; however, a growing body of literature implicates cytokine signaling in the development and/or progression of age-related ocular diseases [50–58]. Indeed, IL-6 and IL-1β inhibitors were therapeutically effective in the context of ocular diseases such as macular edema and ocular surface disease [84–87]. At the same time, however, IL-6 exerts a neuroprotective effect during retinal–RPE separation by promoting photoreceptor survival [88]. Together, these findings underscore the need for future studies to delineate the functional roles of age-related increases in ocular IL-6 and IL-1β and to determine the conditions under which these cytokines contribute to pathology versus protection.

